# Linking structural and functional changes during aging using multilayer brain network analysis

**DOI:** 10.1101/2023.02.15.528643

**Authors:** Gwendolyn Jauny, Mite Mijalkov, Anna Canal-Garcia, Giovanni Volpe, Joana Pereira, Francis Eustache, Thomas Hinault

**Author notes:** Corresponding author: Thomas Hinault, INSERM-EPHE-UNICAEN U1077, 2 rue des Rochambelles, 14032 Caen, FRANCE.

## Abstract

Brain structure and function are intimately linked, however this association remains poorly understood of the complexity of this relationship has remained understudied. Healthy aging is characterized by heterogenous levels of structural integrity changes that influence functional network dynamics. Here, we used the multilayer brain network analysis on structural (diffusion tensor imaging) and functional (magnetoencephalography) data from the Cam-CAN database. We found that the level of similarity of connectivity patterns between brain structure and function in the parietal and temporal regions (alpha frequency band) was associated with cognitive performance in healthy older individuals. These results highlight the impact of structural connectivity changes on the reorganisation of functional connectivity associated with the preservation of cognitive function, and provide a mechanistic understanding of the concepts of brain maintenance and compensation with aging. Investigation of the link between structure and function could thus represent a new marker of individual variability, and of pathological changes.

## Introduction

The brain is one of the most complex biological systems. One of its fascinating aspects, which remains largely unknown, is how wide varieties of brain rhythms and temporally-specific activity patterns^1^ can emerge from a static network architecture^2^. Addressing this issue is a major fundamental endeavor for cognitive neuroscience, which can also improve our understanding of brain changes across the lifespan, and our ability of detecting pathological processes. Previous work has mostly focused on characterizing brain structure (i.e., grey matter and white matter), or brain function (i.e., memory, motor function or cognitive control). These unimodal studies greatly advanced our understating of brain networks and of their associations with cognition^3^. However, brain network analysis methods, such as graph theory, have more recently been applied across modalities to study the interaction between structure and function, showing strong associations between these dimensions^4,5^. Since these seminal studies, the relationship between brain structure and function has been the focus of intense reflection and methodological development, since this relationship is central to many cognitive domains, evolves with age and is affected by pathologies^4^. Here, we investigate these issues in light of age-related brain changes, associated with changes of brain structure that influence neural dynamics^6^, which could further our understanding of the large heterogeneity of individual cognitive trajectories observed during this life period. In particular, structure-function interactions could be central to further understand the preservation (i.e. maintenance^7^ or compensation^8^) or the decline of cognitive performance during aging.

Studying the relationships between white matter fibers (acquired by DTI -diffusion tensor imaging) and Blood-Oxygen-Level-Dependent (BOLD) signal (acquired by fMRI -functional magnetic resonance imaging), previous studies have shown correlations between brain structure and function throughout the lifespan, and particularly across development^9,10^, and during the performance of cognitive tasks^11^. Also, in a healthy older population, Burzynska *et al*.^12^ showed that individuals with preserved white matter fiber integrity had a higher BOLD signal associated with better cognitive performance (see also^13,14^). Many studies have thus focused on this link between structure and function using high-spatial-resolution techniques such as fMRI. However, due to their constrained temporal resolution, age-related changes in the dynamics of the involved networks remain largely understudied.

Previous work has also demonstrated interactions between brain structure and function using high temporal resolution techniques, such as magnetoencephalography (MEG) or electroencephalography (EEG). Indeed, fluctuations in the synchrony and directionality of brain activity have long been considered as noise to be controlled, whereas today they have been reappraised as a fundamental aspect of brain communication^15,16^. These studies have notably highlighted that EEG connectivity is associated with structural connectivity measures in young adults^17^. With healthy aging, Hinault *et al*.^18,19^, showed that a decrease in white matter fiber integrity negatively impacts the neural synchronization between brain regions. However, for all these studies, the interpretation of these interactions is limited as it is based on correlational evidence, which does not account for the full complexity of such a relationship.

A recent approach enables evaluating the relationships between different neuroimaging modalities by constructing a multiplex network model of the brain^20^. This approach allows the creation of a network in which each region is connected to itself across different layers^21^. This technique has already been used in pathology, such as schizophrenia^22^ and Alzheimer’s disease,^23,24^ allowing to highlight brain changes that were not detected in unimodal analyses. Recently, the study by Battiston *et al.^25^*, investigating network connectivity by combining fMRI and DTI data in a two-layer multiplex network revealed relevant relationships between structural and functional brain networks, showing that this technique is an appropriate choice for the study of brain network connectivity. Thus, multiplex brain networks can be used to study the structure-function relationship in healthy aging. To our knowledge, no study has investigated the changes of structural and functional connectivity with increasing age using a multiplex approach applied on DTI and MEG (or EEG) data. However, previous work^26^ suggested that alterations in brain structure can lead to delayed and/or noisier brain communications. Such combination of DTI (structural) and MEG (functional) data in a multiplex connectome in healthy aging is therefore important to identify markers of individual differences and early brain aging effects, preceding major structural changes and loss of functional communications. These changes can lead to deleterious functional consequences^19,27^ or compensatory functional adjustments^28^. This method therefore appears ideal to clarify the association between brain structure and cortical dynamics, to identify the mechanisms underlying cognitive heterogeneity with aging, and the functional adjustments allowing the maintenance of cognitive function.

Here, we propose a multiplex network approach with MEG and DTI data in the context of healthy aging and the associated non-lesional brain changes^29^ (see **Figure 1**). We used the multiplex participation coefficient as an indicator of similarity of connectivity between brain structure and function: a high level of this coefficient corresponded to a similarity of connectivity patterns between these modalities whereas a low level corresponded to a divergence of connectivity patterns between these modalities. We investigated changes in brain structure and function over time in young and older healthy participants from the Cam-CAN database (Cambridge Center for Aging and Neuroscience^30,31^). This database includes multimodal neuroimaging data (MEG, MRI, DTI) as well as cognitive performance evaluation for each individual. Our objectives were two-fold: i) To investigate changes in the interaction between structural integrity levels and synchronized functional networks between young and old individuals, with the underlying hypothesis that a decrease in white matter integrity could negatively impact brain function.. ii) To study the impact of such structure-function relationship on participants’ cognitive performance, where we expected that these changes would be associated with cognitive performance and reveal unique individual differences therein. Compensatory adjustments or maintenance of brain function at the same level as young adults would result in preservation of cognitive performance. Such results could clarify and better characterize maladaptive and compensatory brain communication changes in the presence of aging structural networks.

**Figure 1.**
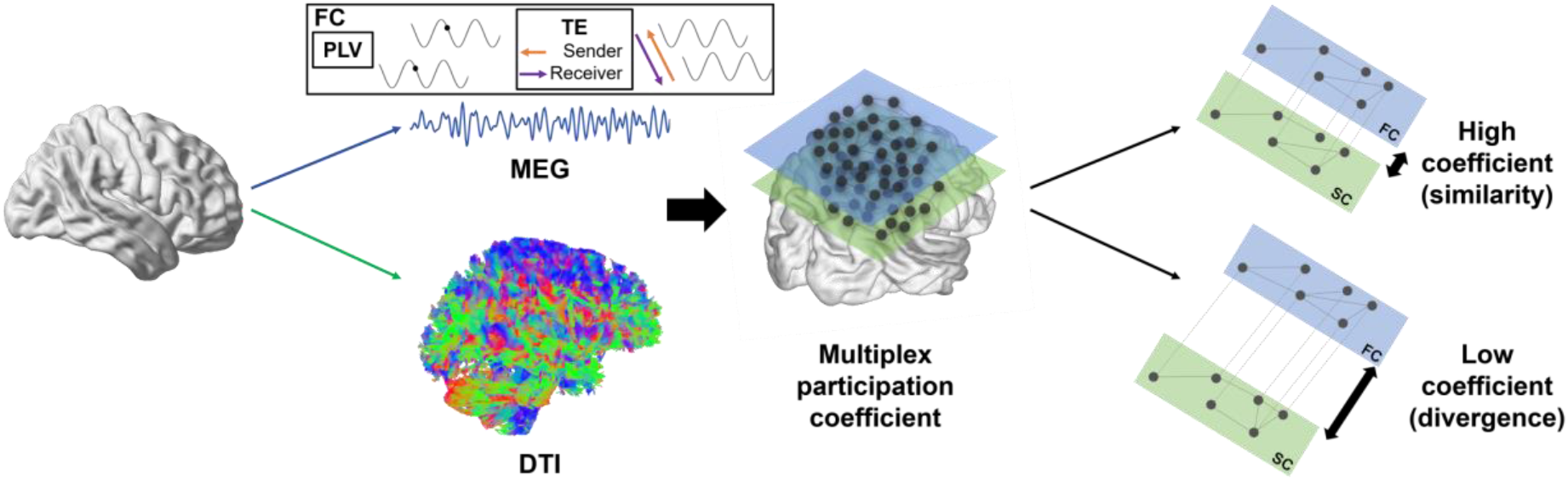
Overview of the creation of the multiplex network from MEG and DTI data. This multiplex network was built with two layers: one representing functional connectivity (FC) from MEG data, either PLV or TE data; the other layer representing structural connectivity (SC) from DTI (anisotropic fraction) data, i.e. FA data. MEG: Magnetoencephalography, DTI: Diffusion Tensor Imaging, PLV: Phase Locking Value, TE: Transfer Entropy, FC: Functional Connectivity, SC: Structural Connectivity

## Results

Two groups of participants (20-30 years for the younger group and 60-70 years for the older group) were formed from the Cam-CAN^30,31^ database. Connectivity analyses were performed on MEG data, and in particular, two measures were studied: phase locking value (PLV), which measure synchrony between regions, and transfer entropy (TE), which measure the directionality of the coupling between brain regions. The data from these two measures were combined with DTI data to form two multiplex structure-function networks (see **Figure 1**). From these networks, the multiplex participation coefficient could be calculated. This coefficient was then studied to determine the level of similarity of connectivity between the two layers (structural and functional) of the network. The different phases of data processing, creation of multiplex networks and statistical analysis are described in the materials and methods section.

### Multiplex network: PLV/DTI

#### Positive association between multiplex participation coefficients and cognitive performance in older adults

Our main objective was to study the effect of healthy aging on structural and functional connectivity, and its association with cognitive abilities (measured with neuropsychological tests assessing working and short-term memory, reasoning ability, executive functions and general cognitive functions, see materials and methods for more information). Thus, we determined which region and which frequency bands age-related changes in multiplex participation coefficient could be associated with cognitive performance. First, we identified the regions and frequency bands that differed between age groups and were associated with cognition: the left temporal and right parietal regions in the alpha frequency band (these two regions showing, respectively, a decrease or an increase in participation in the older individuals compared to the younger). For other regions and frequency bands showing differences not associated with cognitive performance, see **Figure 1s** in supplementary. We found that, for both of these regions, increased multiplex participation coefficient levels were positively associated with cognitive performance in older adults (left temporal/MMSE test, r= 0.313, p= 0.034; right parietal/MMSE test, r= 0.393, p= 0.007; **Figure 2**). No association was found in young adults.

**Figure 2.**
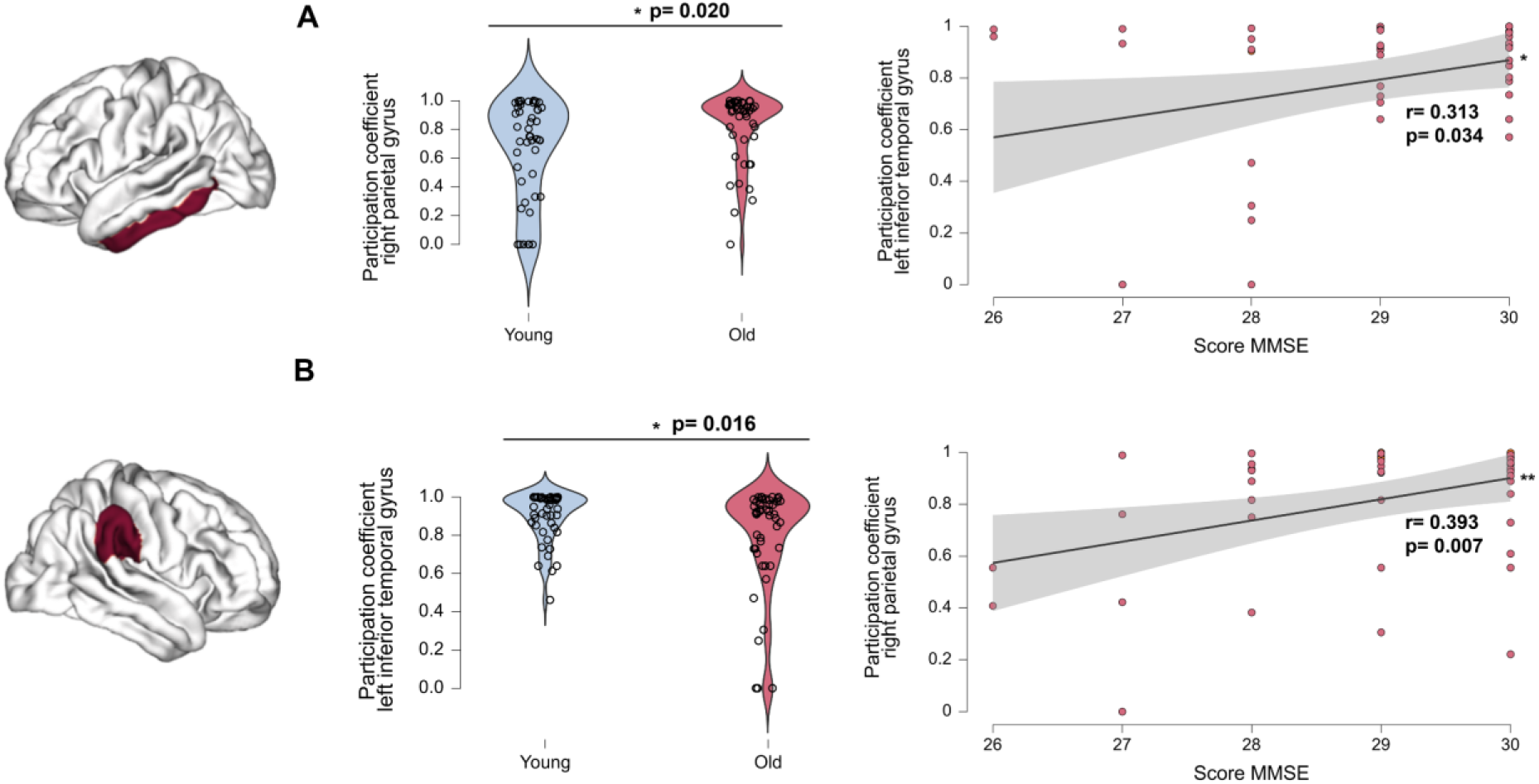
(**A**) Distribution of the young and old groups in left inferior temporal region (t-test) for the multiplex participation coefficient in alpha frequency band for the measure of synchrony (PLV) and positive association between this level of multiplex participation coefficient and MMSE score. (**B**) Distribution of the young and old groups in right parietal region (t-test) for the multiplex participation coefficient in alpha frequency band for the measure of synchrony (PLV) and positive association between this level of multiplex participation coefficient and MMSE score, in older adults. *p<0.05 **p<0.01

#### Maintaining a lower level of multiplex participation coefficient than younger adults is positive for the older population

To further analyze these results, subgroup analyses were performed for these two regions. To do this, participants were grouped according to the level of participation coefficient in each region, forming two groups of older individuals. The older subgroups (i.e., Low participation, High participation; see **Table 1s** to **Table 4s** in supplementary data for the characteristics of each subgroup) did not differ on any measure (e.g., age, gender ratio, level of education, general cognitive performance) other than the level of multiplex participation coefficient (left temporal and right parietal regions). For the left temporal region, young adults differ from both older subgroups, and both subgroups also significantly differ from each other: the level of the participation coefficient was significantly higher for the High participation subgroup than the younger group (p=0.009). The Low participation subgroup showed lower multiplex participation levels than both younger individuals and the High participation subgroup (p<0.001 for both comparison). The Low participation subgroup showed better cognitive performance on the VSTM test than the High participation subgroup (r= 0.584, p= 0.009; **Figure 3A**).

**Figure 3.**
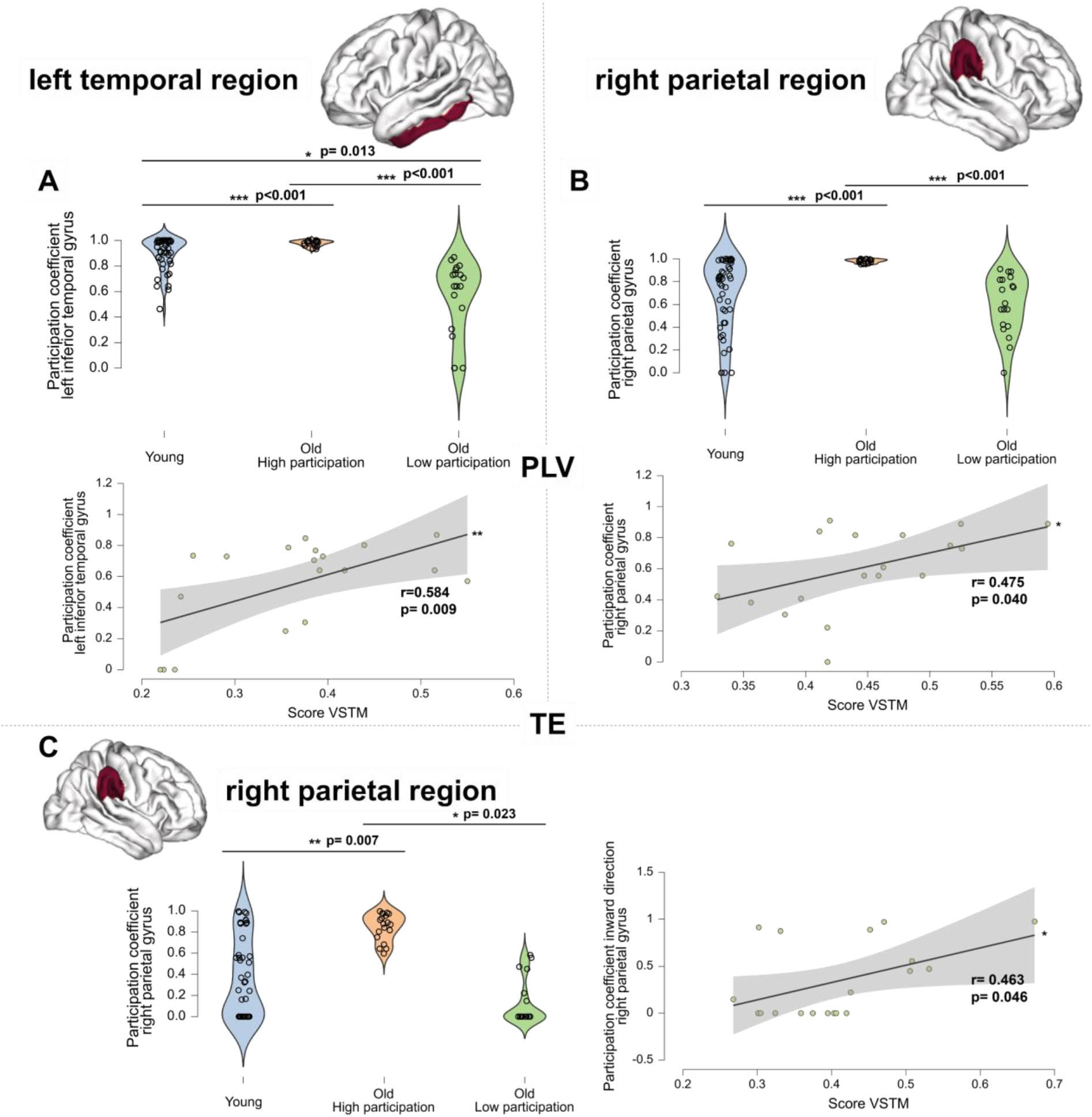
(**A**) Distribution of young adults and older adults’ subgroups for the multiplex participation coefficient in the left temporal region for the measure of synchrony (PLV) in the alpha frequency band. Positive association between participation in the left temporal region and VSTM scores for the Low participation subgroup (regression test; no association with cognition for the High participation older subgroup). (**B**) Distribution of the young adults and older adults’ subgroups for the multiplex participation coefficient in the right parietal region in alpha frequency band. Positive association between participation in the right parietal region and VSTM scores for the Low participation older subgroup (regression test; no association with cognition for the high participation older subgroup). (**C**) Distribution of young adults and older adults’ subgroups for the multiplex participation coefficient in the right parietal region in alpha frequency band for the measure of directionality (TE). Positive association between the participation of the right parietal region and VSTM scores for the Low participation subgroup (regression test; negative association with cognition for the High participation older subgroup: r= −0.491, p=0.033). All results were adjusted for multiple comparisons using FDR corrections at *q* < 0.05. *p<0.05; **p<0.01; ***p<0.001

For the right parietal region, young adults differ from the High participation subgroup, but not with the Low participation subgroup. We observed that the Low participation subgroup (with similar low participation as younger individuals, p=0.962) showed better cognitive performance on the VSTM test (r=0.475, p= 0.040; no association with cognition for the high participation older subgroup; **Figure 3B**).

### Multiplex network TE/DTI

#### Age-related changes in network couplings directionality are positively associated with cognitive performance

Following these results, we examined aging effects and individual differences in these regions using directed functional couplings. For the right parietal region only, in the alpha band, we observed an increase in inward directionality (i.e., directed towards the right parietal region) in older individuals compared to younger individuals (t-test, p=0.038; **Figure 4A**). See Supplementary **Figure 2s** for consistent results involving gamma frequency bands. This increased participation in the inward direction for the right parietal region with aging was positively associated with performance in the VSTM test (r= 0.314, p= 0.034; **Figure 4C**).

**Figure 4.**
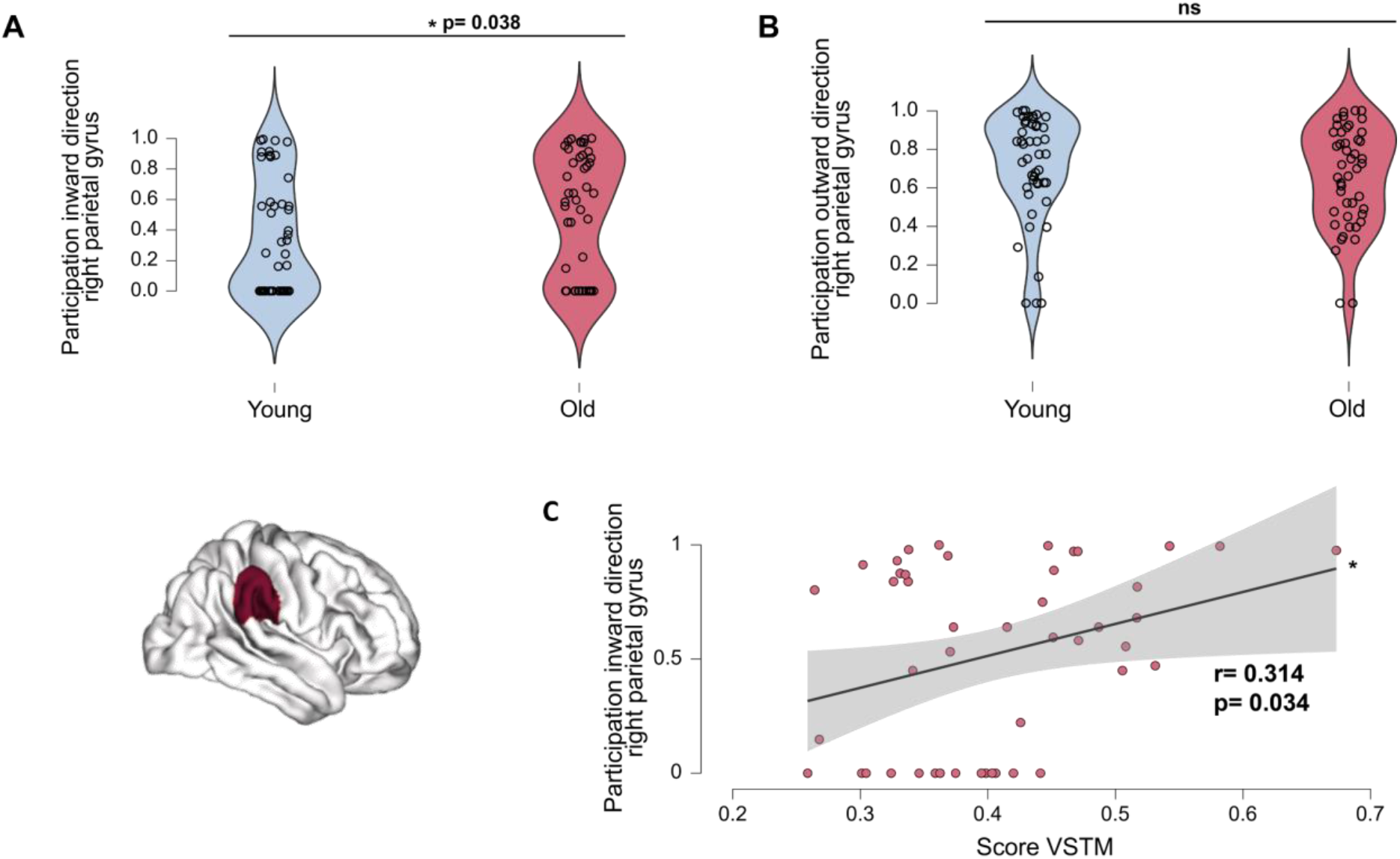
**(A)** Increased inward directionality (i.e., directed towards the right parietal region) in older adults relative to younger adults (t-test) for the right parietal region in alpha frequency band. (**B**) Preserved outward direction (i.e., directed towards other regions of the network) in older adults relative to the younger group for the right parietal region in the alpha frequency band. (**C**) Positive association between the increased multiplex participation coefficient in the inward direction for the right parietal region in alpha frequency band and VSTM test scores (regression test) in the older group. *p<0.05

To further analyze these results, we investigated differences in the same subgroups as in the first part (PLV/DTI) of the results.

We observed that the Low participation subgroup, showing increased inward-directed couplings in right parietal region, also showed better cognitive performance on the VSTM test (r= 0.463, p=0.046; **Figure 3C**) than the High participation subgroup. Supporting these results, the High participation older subgroup showed lower cognitive performance on the VSTM test (r= −0.491, p= 0.033; Figure 3s in supplementary data).

#### Respective contribution of each network layer in younger and older adults

Degree analyses (number of connections) were performed on the respective contribution of each layer, and suggest that the structural layer makes the largest contribution to the reported results, as degree was larger in the structural layer (DTI) than in the functional layer (PLV/TE) for the right parietal region (difference between DTI/PLV and DTI/TE layers, p=.001; see **Figure 3s** in supplementary data). The left temporal region follows this trend as well (difference between DTI/PLV layers, p=0.086; difference between DTI/TE layers, p=0.001).

Interestingly, we examined the contribution of the different layers of connectivity within both older subgroups compared to the younger group for alpha temporal and parietal functional activity (see **Figure 4s).** We observed that the older subgroup that showed lower cognitive performance (High participation) did show difference in contribution between the two functional layers (differences between PLV and TE, p<0.001), in contrast to the Low older subgroup that did show better associations with cognitive performance (p<0.05). These results were found only for the left temporal region.

#### Unique detection of subgroups relative to unimodal network analyses

Finally, we performed unimodal analyses (DTI, MEG) to determine the added value of multiplex analyses relative to functional or structural network investigations. Regarding the structural layer, we replicated the significant difference in white matter integrity between young and old groups (p<0.001) on global connectivity data. Regarding the functional layer, we did not find a significant difference between younger and older adults at the global matrix level, in the alpha frequency band. At the nodal level, no difference between subgroups was observed in functional or structural networks, in contrast with multilayer analyses.

## Discussion

In this study, we have showed the importance of integrating functional and structural information together to better understand aging effects. Our objectives were two-fold: to investigate changes of the brain structure-function association with age, and to determine the impact of changes of this association on cognitive performance in older individuals. Our approach relied on two-layer multiplex network, with a structural layer based on DTI data and another layer based on resting-state MEG data, to identify changes between younger and older healthy individuals from the Cam-CAN repository and to further understand maintenance^7^ and compensation^8^ phenomena observed in aging. Two aspects of functional network connectivity were studied: phase synchrony and directed connectivity. We showed the existence of inter-individual variability at the functional level in older individuals at rest that was associated with cognitive performance. Low structure/function multiplex participation coefficient for structure/synchrony and structure/information transfer in temporal and parietal regions in the alpha frequency band, similar to young adults in parietal region, was associated with preserved cognitive performance in older individuals. These results highlight the impact of fine structural alterations on functional connectivity changes with aging, and provide a better understanding of the relationship between brain structure and function.

The multiplex participation coefficient can be considered as an indicator of co-dependence between modalities: a high level of this coefficient would indicate a high similarity of connectivity between brain structure and function, whereas a low coefficient would indicate a dissociation of structure and function connectivity. Subgroup analyses based on this coefficient allowed the detection of heterogeneity within cognitively healthy older individuals. First, we showed that lower levels of structure/synchrony participation relative to younger adults might be beneficial for cognitive performance. Second, using multiplex structure/directed connectivity network analyses, we showed that low levels of participation in the inward direction (i.e., corresponding to couplings directed towards a given region), to a similar level than young adults, for the regions investigated was beneficial for cognitive performance. In contrast, an increase in this coefficient was found to be negatively associated with cognitive performance. These subgroups were not found in unimodal analyses.

The inferior temporal and supramarginal parietal gyri are both considered to be brain structural cores^32^. They are also both part of the default mode network^33^ (DMN), a network activated at rest, and whose activity has been associated with memory and executive performance^34^. Moreover, the alpha frequency band is involved in the structuring of neural rhythms and has notably been associated with attention allocation and the inhibition of couplings not required for the task^35,36^. By assessing the interaction between brain structure and the alpha frequency band, the present results contribute to existing frameworks about this central brain rhythm^35^, as they did not considered such association. Thus, the disengagement of the DMN, as well as the posterior alpha reduction, are critical for cognition and are impacted by aging^37,38^. Age-related structural changes would be central to these changes and would impact brain function. Our results could indicate that following fine changes in brain architecture, some older individuals will show a lower level of participation coefficient (i.e., a dissociation of connectivity patterns between brain structure and function) than others, which may be due to compensatory functional readjustments involving the alpha frequency band. These changes would enable better cognitive performance than individuals who will not make these functional readjustments, with higher levels of participation coefficient (i.e., a stronger association of connectivity patterns between brain structure and function). Future, longitudinal investigations remain important to further clarify this association.

Our results also reveal that the subgroup of older individuals who showed lower structure/function multiplex participation coefficient, and for whom these changes were positively associated with cognitive performances, showed no difference in contribution (calculated by measuring connectivity levels in each layer) between the phase synchrony (PLV) and information transfer (TE) layers. Conversely, an increase in the contribution of the phase synchrony layer compared to information transfer was found for the group without association with cognition. These results were only observed in the left inferior temporal region. These results could indicate inefficient connectivity in these individuals (i.e., synchronized couplings with little to no information exchange). The observation of synchronized activity may therefore be related to cognitive function, but may also be dissociated from it. Thus, considering synchrony in association with information transfer seems important to clarify age-related changes and to distinguish efficient communications from inefficient/maladaptive network couplings. These communications are highly dependent on the integrity of the underlying structural network, and investigating the respective contribution of structure and function through a multiplex network could also allow distinguishing these functional connectivity patterns in pathologies. Indeed, an increase in neuronal synchrony can be observed in neurodegenerative pathologies and has been considered as maladaptive changes (for a review, see^39^). Further investigations of this distinction could lead to the identification of new markers of subsequent decline and progression of neurodegenerative pathologies.

Several methodological considerations should be discussed regarding the reported results. First, the study of resting-state activity partly limits the direct investigation of the neural bases of cognitive processes, as it might be less directly associated with cognitive functioning than task-related activity^40^. Second, the analysis of layer contributions only showed results for the left inferior temporal region, which does not allow us to generalize our interpretations to the entire brain. Thus, the pattern of layer contributions may be different in other regions and frequency bands^2^, although the reported changes were central in the context of healthy aging. Longitudinal studies could further validate our interpretations and improve our knowledge of other brain regions.

Several questions remain about the association between brain structure and function^6,2^. Indeed, this relationship undergoes crucial changes throughout the lifespan, as well as following several pathologies. The structure-function coupling also appears to fluctuate both over time and regionally. Although structural changes appear to drive changes in coupling between regions, brain functions are not solely determined from brain structure. Decreased integrity impacts neuronal synchrony and information exchange, and these changes are distinctly associated with cognitive performance in individuals. Here, we defined multiplex structure-function models in the context of healthy brain aging to better understand the heterogeneity of these changes across individuals (see **Figure 5** for a schematic representation of this model). In particular, we show its impact on cognitive performance, which improves our knowledge on different theoretical models of aging such as concepts of cognitive maintenance^7^ and compensation^8^. Maintenance would thus be characterized by an imbalance in the contribution of phase synchrony and transfer information: with a higher level of contribution from PLV than from TE. Moreover, the level of similarity of connectivity between brain structure and function would be very low. Cognitive decline would also be associated with an imbalance in the contribution of phase synchrony and transfer information. However, in contrast to maintenance, the level of similarity of connectivity between brain structure and function would be very high. Finally, Compensation would be characterized by a balance in the contribution of phase synchrony and transfer information. The level of similarity of connectivity between structure and brain function would be very low, in the same way as in the maintenance concept. Indeed, a dissociation of connectivity pattern between structure and function has been associated with the preservation of cognitive performance. Importantly, these individual markers were not found in unimodal analyses. This new approach might yield a better understanding of the brain, which could be useful in clinical applications to better understand certain pathologies such as neurodegenerative diseases, and more generally to further our understanding of the link between structure and function in the brain.

**Figure 5.**
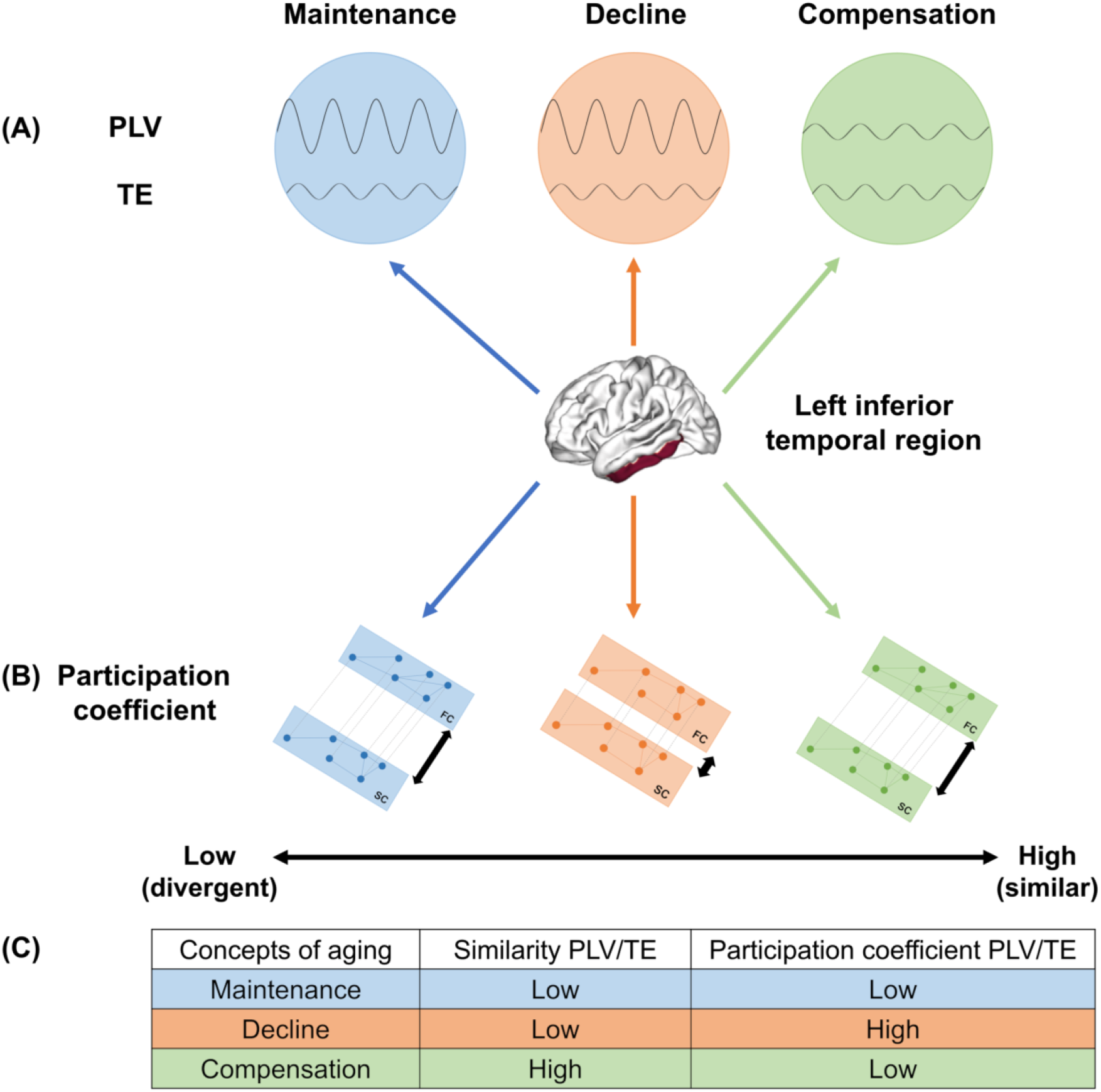
Schematic representation of the proposed model for the left inferior temporal region. (A) Level of contribution for PLV and TE. (B) Participation coefficient for PLV/DTI and TE/DTI multiplex network. (C) Summary of the relation between the level of similarity of contribution from PLV/TE, participation coefficient and concepts of aging. DTI: Diffusion Tensor Imaging, PLV: Phase Locking Value, TE: Transfer Entropy, FC: Functional Connectivity, SC: Structural Connectivity

## Materials and Methods

### Participants

All participants aged 20-30 years and 60-70 years were selected from the Cam-CAN database^30,31^, in line with demographic characteristics of individuals recruited in previous work^41,18^. Thus, we analysed data from 46 young (29 women and 17 men; aged 22-29 years) and 46 older healthy adults (29 women and 17 men; aged 60-69 years) (**Table 1**). All participants were right-handed, showed normal cognitive functioning^42^ (Montreal Cognitive Assessment (MoCA) score >26), and no neurological or psychiatric conditions.

**Table 1.** Demographics and scores for both groups younger and older participants. MMSE: Mini-Mental State Evaluation; VSTM: Visual Short-Term Memory; Hotel_num_rows: corresponding to the number of rows performed by the participant; Hotel_Time: corresponding to the time used to performed all rows by the participant. Differences between the two groups were calculated using t-test.

| Variables | Young adults | Older adults | p-value |
| --- | --- | --- | --- |
| Number of participants | 46 | 46 | 1.000 |
| Number of women | 29 | 29 | 1.000 |
| Age | 26.5 (2.01) | 64.5 (2.85) | **0.001** |
| **Years of education** | **22.2** (2.873) | **19.1** (3.262) | **0.001** |
| **MMSE** | **29.5** (0.863) | **28.9** (1.173) | **0.013** |
| **VSTM** | **0.5** (0.088) | **0.4** (0.069) | **0.001** |
| **Cattell** | **37.8** (3.628) | **30.5** (6.285) | **0.001** |
| **Hotel_Num_rows** | **4.7** (0.585) | **4.3** (1.008) | **0.018** |
| **Hotel_Time** | **227.7** (119.796) | **326.9** (194.305) | **0.005** |

### Behavioural measures

A detailed description of behavioural measures can be found in supplementary materials (see also Refs. ^30,31^). Cognitive performance was assessed with the Mini-Mental State Evaluation^43^ (MMSE) as a measure of general cognitive functioning, the Visual Short-Term Memory^44^ (VSTM) as a test of short-term memory and working-memory maintenance, the Cattel test^45^ measuring reasoning ability, and the Hotel Test^46^ assessing executive functions (notably planning abilities). Despite significant differences between the two groups, all participants had normal cognitive function. These variables were added as covariates in statistical analyses.

### MEG, structural MRI and DTI data acquisition

Resting MEG activity was measured for 10 minutes (sampling rate: 1kHz, bandpass filter: 0.03-330 Hz) with a 306-channel MEG system. Participants’ 3D-T1 MRI images were acquired on a 32-channel 3T MRI scanner. The following parameters were used: repetition time = 2250 ms; echo time = 2.99 ms; inversion time = 900 ms; flip angle = 9 degrees; field of view = 256 mm x 240 mm x 192 mm; voxel size = 2 mm isotropic; GRAPPA acceleration factor = 2; acquisition time = 4 minutes and 32 seconds. DTI data were obtained with the following parameters: repetition time = 9100 ms; echo time = 104 ms; inversion time = 900 ms; field of view = 192 mm x 192 mm; 66 axial slices; voxel size = 2 mm isotropic; B0 = 0.1000/2000s/mm2; acquisition time = 10 minutes and 2 seconds, readout time 0.0684 (echo spacing = 0.72ms, EPI factor = 96). See https://camcan-archive.mrc-cbu.cam.ac.uk/dataaccess/ for more information.

### MEG data pre-processing

The Elekta Neuromag MaxFilter 2.2 has been applied to MEG data (temporal signal space separation (tSSS): 0.98 correlation, 10s window; bad channel correction: ON; motion correction: OFF; 50Hz+harmonics (mains) notch). Afterwards, artifact rejection, filtering (0.3-100 Hz bandpass), temporal segmentation into epochs, averaging and source estimation were performed using Brainstorm^47^. In addition, physiological artefacts (e.g., blinks, saccades) were identified and removed using spatial space projection of the signal. In order to improve the accuracy of the source reconstruction, the FreeSurfer^48^ software was used to generate cortical surfaces and automatically segment them from the cortical structures from each participant’s T1-weighted anatomical MRI. The advanced MEG model was obtained from a symmetric boundary element method (BEM model^49^; OpenMEEG^50^), fitted to the spatial positions of each sensor^51^. A cortically constrained sLORETA procedure was applied to estimate the cortical origin of the scalp MEG signals. The estimated sources were then smoothed and projected into standard space (i.e., ICBM152 template) for comparisons between groups and individuals, while accounting for differences in anatomy (i.e., gray matter). This procedure was applied for the entire recording duration.

### Connectivity analyses

Phase-locking value analyses^52^ (PLV) were used to determine the functional synchrony between regions of interest. PLV estimates the variability of phase differences between two regions over time. If the phase difference varies little, the PLV is close to 1 (this corresponds to high synchronisation between the regions), while the low association of phase difference across regions is indicated by a PLV value close to zero. To ensure PLV results did not reflect volume conduction artefacts, additional control analyses were conducted using phase lag index (weighted PLI analyses). Because PLV is an undirected measure of functional connectivity, and to investigate brain dynamics with complementary metrics, analyses of transfer entropy (TE) have also been conducted. TE measures how a signal can predict subsequent changes in another signal^53^. It thus provides a directed measure of a coupling’s strength. If there is no coupling between regions, then TE is close to 0, while TE is close to 1 if there is a strong coupling between two regions.

The range of each frequency band was based on the frequency of the individually observed alpha peak frequency (IAF), measured as the average of peaks detected from both occipitoparietal magnetometers and gradiometers. In line with previous work^54^ the following frequency bands were considered: Delta (IAF-8/IAF-6), Theta (IAF-6/IAF-2), Alpha (IAF-2/IAF+2), Beta (IAF+2/IAF+14), Gamma (IAF+15/IAF+80). To reduce the dimensionality of the data, the first component of the principal component analysis (PCA) decomposition of the time course of activation in each of the 68 regions of interest (ROI) from the Desikan-Killiany brain atlas. The first component, rather than the average activity, was chosen to reduce signal leakage^55^.

### DTI data pre-processing

Pre-processing of the diffusion data was performed using ExploreDTI^56^ and included the following steps: (a) images were corrected for eddy current distortions and participant motion; (b) a non-linear least squares method was applied for diffusion tensor estimation, and (c) deterministic DTI tractography was applied using the following parameters: uniform resolution of 2 mm, fractional anisotropy (FA) threshold of 0.2 (limit: 1), angle threshold of 45°, and fibre length range of 50 to 500 mm. The network analysis tools in ExploreDTI were used to quantify the FA value of the fibres connecting the regions of the Desikan atlas, to obtain similar matrices to MEG data, using Freesurfer’s individual cortical parcellation.

### Multiplex Network construction and measures

Using BRAPH^57^ software (http://braph.org/), a multiplex network was defined for each subject, with two layers: one “structural” layer with DTI tract FA data, and one “functional” layer with PLV or TE MEG data (in this study, a simplification of TE was used to determine whether a region was a receiver or sender). TE analyses were performed on each region and distinguished coupling directed from the network towards a given region (i.e., the inward direction), or from a given region towards the rest of the network (i.e., the outward direction). In each layer, brain regions from the Desikan-Killiany atlas^58^ are represented by nodes connected by edges (see a method’s summary in **Figure 1**). A binary multiplex matrix was calculated from the individual matrices of DTI and MEG data of each participant. Auto-correlations between regions were excluded from the analyses.

To evaluate across-layer integration, the multiplex participation coefficient^59^ was investigated, allowing the quantification of the connectivity similarity of a node across the different layers. The multiplex participation coefficient of a node *i* is defined as^59^: 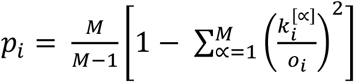 where M is the number of layers, 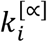 the degree of node *i* at the ∝ –*th* layer and *o_i_* is the overlapping degree of node *i*, 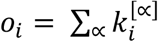. This coefficient measures how similar the connectivity patterns are in both layers of the multiplex network. Values range between 0 and 1. In particular, value of 1 means that the node makes the exact connections in both layers, while a value of 0 means that the nodes connections in both layers are different from each other. A large participation value indicates that the node may be central or a hub. To determine which layer is driving the observed results, the degree (i.e., number of connections of each layer of the multiplex network for a given region) was also calculated for each group as: 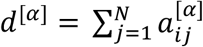; where 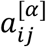 is the link between node i and j in layer *α*.

### Statistical tests

To assess differences between age groups in multiplex participation for different brain regions, t-tests were applied using the Jamovi software (https://www.jamovi.org/; version 1.6.23). Regression analyses were performed in the older adults’ group to assess whether the level of participation coefficient for a region was associated with cognitive performance. Afterwards, participants were grouped according to the level of participation coefficient for each region. Two subgroups were then formed: one corresponding to individuals with a high participation coefficient called “High participant group” and another with a low participation coefficient called “Low participant group”. The median individuals (four from each group) were removed from subgroup analyses to reduce median split bias. As a result, each subgroup was composed of 19 individuals. Subgroups were also found in young adults but due to the large variability in young individuals, were considered as a single group. T-tests were also performed to determine differences between subgroups. Original degrees of freedom and corrected p-values are reported. Results were FDR corrected for multiple comparisons^60^.

## Acknowledgements

This research did not receive any specific grant from funding agencies in the public, commercial, or not-for-profit sectors.

## Author Contributions

G.Jauny: Investigation, Analysis, Writing; M.Mijalkov: Methodology, Software, Review; A.Canal-Garcia: Methodology, Software, Review; G.Volpe: Methodology, Software, Review; J.B.Pereira: Methodology, Software, Review; F.Eustache: Supervision, Review; T.Hinault: Conceptualization, Methodology, Supervision, Review.

## References

1. Buzsáki, G. Rhythms of the Brain. (Oxford University Press, 2006). doi:10.1093/acprof:oso/9780195301069.001.0001.

2. Liu, Z.-Q. et al. Time-resolved structure-function coupling in brain networks. Commun. Biol. 5, 1–10 (2022).

3. Park, H.-J. & Friston, K. Structural and functional brain networks: from connections to cognition. Science 342, 1238411 (2013).

4. Bullmore, E. & Sporns, O. Complex brain networks: graph theoretical analysis of structural and functional systems. Nat. Rev. Neurosci. 10, 186–198 (2009).

5. Honey, C. J. et al. Predicting human resting-state functional connectivity from structural connectivity. Proc. Natl. Acad. Sci. 106, 2035–2040 (2009).

6. Suárez, L. E., Markello, R. D., Betzel, R. F. & Misic, B. Linking Structure and Function in Macroscale Brain Networks. Trends Cogn. Sci. 24, 302–315 (2020).

7. Nyberg, L., Lövdén, M., Riklund, K., Lindenberger, U. & Bäckman, L. Memory aging and brain maintenance. Trends Cogn. Sci. 16, 292–305 (2012).

8. Cabeza, R. et al. Maintenance, reserve and compensation: the cognitive neuroscience of healthy ageing. Nat. Rev. Neurosci. 2018 1911 19, 701–710 (2018).

9. Uddin, L. Q., Supekar, K. S., Ryali, S. & Menon, V. Dynamic reconfiguration of structural and functional connectivity across core neurocognitive brain networks with development. J. Neurosci. Off. J. Soc. Neurosci. 31, 18578–18589 (2011).

10. Baum, G. L. et al. Development of structure–function coupling in human brain networks during youth. Proc. Natl. Acad. Sci. 117, 771–778 (2020).

11. Medaglia, J. D. et al. Functional alignment with anatomical networks is associated with cognitive flexibility. Nat. Hum. Behav. 2, 156–164 (2018).

12. Burzynska, A. Z. et al. White Matter Integrity Supports BOLD Signal Variability and Cognitive Performance in the Aging Human Brain. PLOS ONE 10, e0120315 (2015).

13. Webb, C. E., Rodrigue, K. M., Hoagey, D. A., Foster, C. M. & Kennedy, K. M. Contributions of White Matter Connectivity and BOLD Modulation to Cognitive Aging: A Lifespan Structure-Function Association Study. Cereb. Cortex 30, 1649–1661 (2020).

14. Hinault, T., Larcher, K., Bherer, L., Courtney, S. M. & Dagher, A. Age-related differences in the structural and effective connectivity of cognitive control: a combined fMRI and DTI study of mental arithmetic. Neurobiol. Aging 82, 30–39 (2019).

15. Uddin, L. Q. Bring the Noise: Reconceptualizing Spontaneous Neural Activity. Trends Cogn. Sci. 24, 734–746 (2020).

16. Untergehrer, G., Jordan, D., Kochs, E. F., Ilg, R. & Schneider, G. Fronto-Parietal Connectivity Is a Non-Static Phenomenon with Characteristic Changes during Unconsciousness. PLOS ONE 9, e87498 (2014).

17. Deslauriers-Gauthier, S. et al. White matter information flow mapping from diffusion MRI and EEG. NeuroImage 201, 116017 (2019).

18. Hinault, T., Kraut, M., Bakker, A., Dagher, A. & Courtney, S. M. Disrupted neural synchrony mediates the relationship between white matter integrity and cognitive performance in older adults. Cereb. Cortex 30, 5570–5582 (2020).

19. Hinault, T. et al. Age-related differences in network structure and dynamic synchrony of cognitive control. NeuroImage 236, 118070 (2021).

20. Vaiana, M. & Muldoon, S. F. Multilayer Brain Networks. J. Nonlinear Sci. 30, 2147–2169 (2020).

21. Battiston, F., Guillon, J., Chavez, M., Latora, V. & De Vico Fallani, F. Multiplex core–periphery organization of the human connectome. J. R. Soc. Interface 15, 20180514 (2018).

22. Brookes, M. J. et al. A multi-layer network approach to MEG connectivity analysis. NeuroImage 132, 425–438 (2016).

23. Canal-Garcia, A. et al. Multiplex connectome changes across the alzheimer’s disease spectrum using gray matter and amyloid data. Cereb. Cortex 32, 3501–3515 (2022).

24. Guillon, J. et al. Loss of brain inter-frequency hubs in Alzheimer’s disease. Sci. Rep. 7, 10879 (2017).

25. Battiston, F., Nicosia, V., Chavez, M. & Latora, V. Multilayer motif analysis of brain networks. Chaos Interdiscip. J. Nonlinear Sci. 27, 047404 (2017).

26. Courtney, S. M. & Hinault, T. When the time is right: Temporal dynamics of brain activity in healthy aging and dementia. Prog. Neurobiol. 203, 102076 (2021).

27. Tóth, B. et al. Frontal midline theta connectivity is related to efficiency of WM maintenance and is affected by aging. Neurobiol. Learn. Mem. 114, 58–69 (2014).

28. Ariza, P. et al. Evaluating the effect of aging on interference resolution with time-varying complex networks analysis. Front. Hum. Neurosci. 9, (2015).

29. Park, D. C. & Reuter-Lorenz, P. The Adaptive Brain: Aging and Neurocognitive Scaffolding. Annu. Rev. Psychol. 60, 173 (2009).

30. Shafto, M. A. et al. The Cambridge Centre for Ageing and Neuroscience (Cam-CAN) study protocol: a cross-sectional, lifespan, multidisciplinary examination of healthy cognitive ageing. BMC Neurol. 14, (2014).

31. Taylor, J. R. et al. The Cambridge Centre for Ageing and Neuroscience (Cam-CAN) data repository: Structural and functional MRI, MEG, and cognitive data from a cross-sectional adult lifespan sample. NeuroImage 144, 262–269 (2017).

32. Hagmann, P. et al. Mapping the Structural Core of Human Cerebral Cortex. PLOS Biol. 6, e159 (2008).

33. Andrews-Hanna, J. R., Smallwood, J. & Spreng, R. N. The default network and self-generated thought: component processes, dynamic control, and clinical relevance. Ann. N. Y. Acad. Sci. 1316, 29–52 (2014).

34. Andrews-Hanna, J. R. et al. Disruption of Large-Scale Brain Systems in Advanced Aging. Neuron 56, 924–935 (2007).

35. Bonnefond, M., Kastner, S. & Jensen, O. Communication between Brain Areas Based on Nested Oscillations. eNeuro 4, ENEURO.0153-16.2017 (2017).

36. Sadaghiani, S. & Kleinschmidt, A. Brain Networks and α-Oscillations: Structural and Functional Foundations of Cognitive Control. Trends Cogn. Sci. 20, 805–817 (2016).

37. Anderson, B. A., Folk, C. L. & Courtney, S. M. Neural mechanisms of goal-contingent task disengagement: Response-irrelevant stimuli activate the default mode network. Cortex 81, 221–230 (2016).

38. Poza, J. et al. Phase-amplitude coupling analysis of spontaneous EEG activity in Alzheimer’s disease. Annu. Int. Conf. IEEE Eng. Med. Biol. Soc. IEEE Eng. Med. Biol. Soc. Annu. Int. Conf. 2017, 2259–2262 (2017).

39. Jauny, G., Eustache, F. & Hinault, T. T. M/EEG Dynamics Underlying Reserve, Resilience, and Maintenance in Aging: A Review. Front. Psychol. 13, (2022).

40. Grigg, O. & Grady, C. L. Task-Related Effects on the Temporal and Spatial Dynamics of Resting-State Functional Connectivity in the Default Network. PLOS ONE 5, e13311 (2010).

41. Coquelet, N. et al. The electrophysiological connectome is maintained in healthy elders: A power envelope correlation MEG study. Sci. Rep. 7, 1–10 (2017).

42. Nasreddine, Z. S. et al. The Montreal Cognitive Assessment, MoCA: A Brief Screening Tool For Mild Cognitive Impairment. J. Am. Geriatr. Soc. 53, 695–699 (2005).

43. Folstein, M. F., Folstein, S. E. & McHugh, P. R. “Mini-mental state”: A practical method for grading the cognitive state of patients for the clinician. J. Psychiatr. Res. 12, 189–198 (1975).

44. Vogel, E. K., Woodman, G. F. & Luck, S. J. Storage of features, conjunctions, and objects in visual working memory. J. Exp. Psychol. Hum. Percept. Perform. 27, 92–114 (2001).

45. Horn, J. L. & Cattell, R. B. Refinement and test of the theory of fluid and crystallized general intelligences. J. Educ. Psychol. 57, 253–270 (1966).

46. Shallice, T. & Burgess, P. W. DEFICITS IN STRATEGY APPLICATION FOLLOWING FRONTAL LOBE DAMAGE IN MAN. Brain 114, 727–741 (1991).

47. Tadel, F., Baillet, S., Mosher, J. C., Pantazis, D. & Leahy, R. M. Brainstorm: A user-friendly application for MEG/EEG analysis. Comput. Intell. Neurosci. 2011, (2011).

48. Fischl, B. FreeSurfer. NeuroImage 62, 774–781 (2012).

49. Kybic, J. et al. A common formalism for the Integral formulations of the forward EEG problem. IEEE Trans. Med. Imaging 24, 12–28 (2005).

50. Gramfort, A., Papadopoulo, T., Olivi, E. & Clerc, M. OpenMEEG: opensource software for quasistatic bioelectromagnetics. Biomed. Eng. OnLine 9, 45 (2010).

51. Huang, M. X., Mosher, J. C. & Leahy, R. M. A sensor-weighted overlapping-sphere head model and exhaustive head model comparison for MEG. Phys. Med. Biol. 44, 423 (1999).

52. Lachaux, J.-P., Rodriguez, E., Martinerie, J. & Varela, F. J. Measuring Phase Synchrony in Brain Signals. Hum Brain Mapp. 8, 194–208 (1999).

53. Ursino, M., Ricci, G. & Magosso, E. Transfer Entropy as a Measure of Brain Connectivity: A Critical Analysis With the Help of Neural Mass Models. Front. Comput. Neurosci. 14, 45 (2020).

54. Toppi, J. et al. Different Topological Properties of EEG-Derived Networks Describe Working Memory Phases as Revealed by Graph Theoretical Analysis. Front. Hum. Neurosci. 11, 637 (2017).

55. Sato, M., Yamashita, O., Sato, M. aki & Miyawaki, Y. Information spreading by a combination of MEG source estimation and multivariate pattern classification. PLOS ONE 13, e0198806 (2018).

56. Leemans, A., Jeurissen, B., Sijbers, J. & Jones, D. K. ExploreDTI: a graphical toolbox for processing, analyzing, and visualizing diffusion MR data.

57. Mijalkov, M. et al. BRAPH: A graph theory software for the analysis of brain connectivity. PLOS ONE 12, e0178798 (2017).

58. Desikan, R. S. et al. An automated labeling system for subdividing the human cerebral cortex on MRI scans into gyral based regions of interest. NeuroImage 31, 968–980 (2006).

59. Battiston, F., Nicosia, V. & Latora, V. Structural measures for multiplex networks. Phys. Rev. E 89, 032804 (2014).

60. Benjaminit, Y. & Hochberg, Y. Controlling the False Discovery Rate: a Practical and Powerful Approach to Multiple Testing. J R Stat. Soc B 57, 289–300 (1995).

