## Supplementary data for "Linking structural and functional changes during aging using multilayer brain network analysis"

Complete characterization of the cognitive tests. These descriptions are based on what can be found on the site [https://www.cam-can.org/](https://www.cam-can.org/) and in Shafto *et al.* (2014) and Taylor *et al.* (2017).

**MMSE** (Mini-Mental State Examination): Test comprising a series of 30 questions divided into 6 categories: evaluation of spatial and temporal orientation abilities, learning abilities, attention and calculation abilities, information recall abilities, language and praxis abilities.

**VSTM**: View (1–4) coloured discs briefly presented on a computer screen, then after a delay, attempt to remember the colour of the disc that was at a cued location, with response indicated by selecting the colour on a colour wheel (touchscreen input).

**Cattell**: The test contains four subtests with different types of nonverbal "puzzles": series completion, classification, matrices, and conditions. Correct responses are given a score of 1 for a total maximum score of 46.

**Hotel task**: Perform tasks in role of hotel manager: write customer bills, sort money, proofread advert, sort playing cards, alphabetise list of names. Total time must be allocated equally between tasks; there is not enough time to complete any one task. The number of actions performed (Hotel_num) and the time taken (Hotel_time) are measured.

**Figure 1s: Description of regions with statistically significant differences in multiplex participation coefficients between young and old groups**

**A**

| Regions | Participation Young group | Participation Old group | p-value | Effect | Frequency bands |
|---|---|---|---|---|---|
| **Inferiortemporal L** | 0.901 (0.129) | 0.790 (0.277) | ≤ 0.041 | **Smaller** participation in older adults | Alpha, Beta, Theta |
| Rostralmiddlefrontal L | 0.840 (0.168) | 0.726 (0.336) | ≤ 0.042 | **Smaller** participation in older adults | Delta, Gamma |
| Enthorinal L | 0.686 (0.318) | 0.821 (0.272) | ≤ 0.031 | **Bigger** participation in older adults | Alpha, Delta |
| **Supramarginal R** | 0.668 (0.343) | 0.815 (0.245) | ≤ 0.029 | **Bigger** participation in older adults | Alpha, Delta, Theta |
| Insula R | 0.473 (0.383) | 0.635 (0.365) | 0.040 | **Bigger** participation in older adults | Theta |

**B**

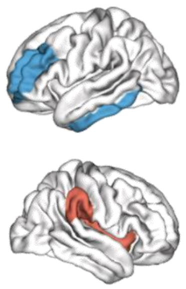

**Figure 1s**. (**A**) Table of the five brain regions where multiplex participation coefficient was found to differ between the two groups, young and old (t-test), with the two regions in bold showing significant associations with behavioral performance. (**B**) Visualization of the five brain regions. In blue, regions where multiplex participation coefficient was decreased; in green, regions where multiplex participation coefficient was increased in the older group compared to the younger group. All results were adjusted for multiple comparisons using FDR corrections at $q < 0.05$.

**Table 1s. Demographics and scores for younger group and older subgroups participants formed from multiplex participation coefficient in left temporal region.**

| Variables | Young adults | Older adults | p-value |
|---|---|---|---|

| | | High participation | Low participation | - |
|---|---|---|---|---|
| Number of participants | 46 | 19 | 19 | - |
| Number of females | 29 | 9 | 10 | - |
| **Age** | **26.5 (2.01)** | **64.4 (2.98)** | **64.5 (2.84)** | **< 0.001** |
| Years of education | 22.2 (2.87) | 20.1 (3.3) | 19.2 (3.2) | **< 0.015** |
| MMSE | 29.5 (0.863) | 28.8 (1.4) | 29 (1.05) | **0.041**/0.051 |
| VSTM_all | 0.5 (0.0875) | 0.430 (0.057) | 0.430 (0.081) | **<0.004** |
| Cattell | 37.8 (3.63) | 30.84 (6.89) | 30.57 (6.50) | **<0.001** |
| Hotel_Num_rows | 4.7 (0.585) | 4.21 (0.91) | 4.36 (1.16) | **0.009**/0.108 |
| Hotel_Time | 227.7 (120) | 332.2 (181.9) | 334.7 (210.3) | **< 0.013** |
| **Inferiortemporal_participation PLV** | **0.902 (0.125)** | **0.977 (0.0217)** | **0.552 (0.295)** | **< 0.013** |
| **Inferiortemporal_participation TE** | **0.379 (0.37)** | **0.784 (0.25)** | **0.494 (0.32)** | **< 0.001**/0.244 |

The p-values correspond to the highest p-value between the younger group and both older subgroups when older subgroups differ. In bold: p-value statistically different. In green: p-value between the younger and the older Low participation groups when older subgroups differ.

**Table 2s. Demographics and scores for both older subgroups participants formed from multiplex participation coefficient in left temporal region.**

| Variables | Older adults | | p-value |
|---|---|---|---|
| | High participation | Low participation | - |
| Number of participants | 19 | 19 | - |
| Number of females | 9 | 10 | - |
| Age | 64.4 (2.98) | 64.5 (2.84) | 0.960 |
| Years of education | 20.1 (3.3) | 19.2 (3.2) | 0.455 |
| MMSE | 28.8 (1.4) | 29 (1.05) | 0.799 |
| VSTM_all | 0.430 (0.057) | 0.430 (0.081) | 0.992 |
| Cattell | 30.84 (6.89) | 30.57 (6.50) | 0.904 |
| Hotel_Num_rows | 4.21 (0.91) | 4.36 (1.16) | 0.645 |
| Hotel_Time | 332.2 (181.9) | 334.7 (210.3) | 0.970 |
| **Inferiortemporal_participation PLV** | **0.977 (0.0217)** | **0.552 (0.295)** | **<0.001** |

| **Inferiortemporal_participation TE** | **0.784 (0.25)** | **0.494 (0.32)** | **0.004** |
|---|---|---|---|

In bold: p-value statistically different between both older subgroups.

**Table 3s. Demographics and scores for younger group and older subgroups participants formed from multiplex participation coefficient in right parietal region.**

| Variables | Young adults | Older adults | | p-value |
|---|---|---|---|---|
| | | High participation | Low participation | - |
| Number of participants | 46 | 23 | 23 | - |
| Number of females | 29 | 8 | 11 | - |
| **Age** | **26.5 (2.01)** | **63.913 (2.85)** | **65.087 (2.84)** | **< 0.001** |
| Years of education | 22.2 (2.87) | 19.3 (2.81) | 19 (2.34) | **< 0.001** |
| MMSE | 29.5 (0.863) | **29.31 (0.88)** | **28.52 (1.42)** | 0.440 / **0.001** |
| VSTM_all | 0.5 (0.0875) | 0.428 (0.070) | 0.443 (0.069) | **<0.014** |
| Cattell | 37.8 (3.63) | 29.78 (5.23) | 31.47 (6.89) | **<0.001** |
| Hotel_Num_rows | 4.7 (0.5852) | 4.452 (0.69) | 4.21 (1.27) | 0.242 / **0.030** |
| Hotel_Time | 227.7 (119.7) | 281.5 (144.14) | 347.5 (232.05) | 0.129/**0.009** |
| **Supramarginal_participation PLV** | **0.668 (0.3205)** | **0.981 (0.016)** | **0.601 (0.25)** | **<0.001 /** 0.447 |
| **Supramarginal_participation TE** | **0.351 (0.37)** | **0.634 (0.36)** | **0.341 (0.40)** | **0.007**/0.923 |

The p-values correspond to the highest p-value between the younger group and both older subgroups when older subgroups differ. In bold: p-value statistically different. In red: p-value between the younger and the older High participation groups when older subgroups differ. In green: p-value between the younger and the older Low participation groups when older subgroups differ.

**Table 4s. Demographics and scores for both older subgroups participants formed from multiplex participation coefficient in right parietal region.**

| Variables | Older adults | | p-value |
|---|---|---|---|
| | High participation | Low participation | - |
| Number of participants | 23 | 23 | - |
| Number of females | 8 | 11 | - |

| | | | |
|---|---|---|---|
| Age | 63.913 (2.85) | 65.087 (2.84) | 0.244 |
| Years of education | 19.3 (2.81) | 19 (2.34) | 0.718 |
| **MMSE** | **29.31 (0.88)** | **28.52 (1.42)** | **0.048** |
| VSTM_all | 0.428 (0.070) | 0.443 (0.069) | 0.521 |
| Cattell | 29.78 (5.23) | 31.47 (6.89) | 0.402 |
| Hotel_Num_rows | 4.452 (0.69) | 4.21 (1.27) | 0.349 |
| Hotel_Time | 281.5 (144.14) | 347.5 (232.05) | 0.300 |
| **Supramarginal_participation PLV** | **0.981 (0.016)** | **0.601 (0.25)** | **<0.001** |
| **Supramarginal_participation TE** | **0.634 (0.36)** | **0.341 (0.40)** | **0.023** |

In bold: p-value statistically different between both older subgroups.

**Figure 2s: Changing network directed couplings in left temporal region with age is cognitively beneficial for older adults**

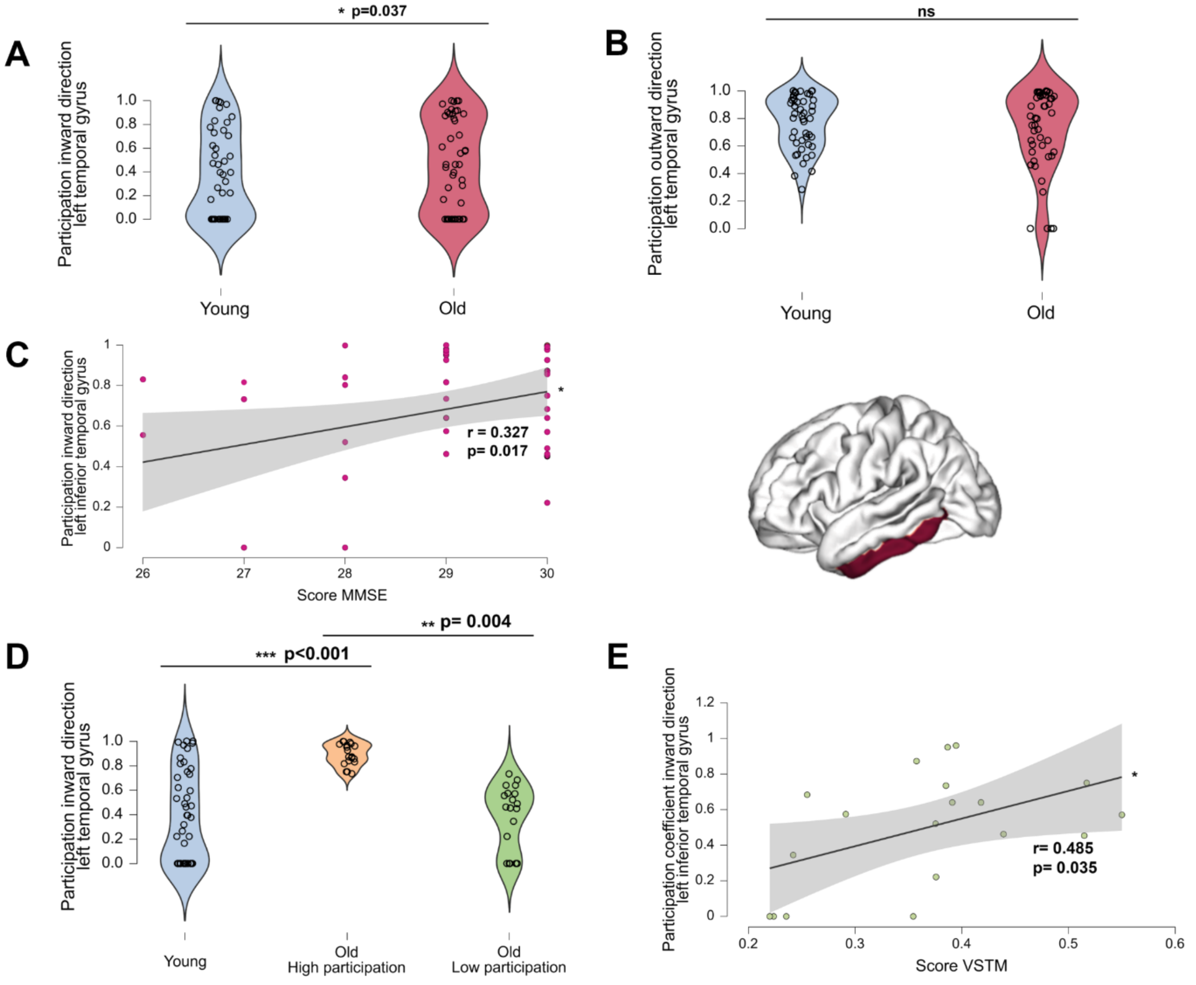

**Figure 2s**. (**A**) Increased inward directionality (i.e., directed towards the right parietal region) in the older group compared to the younger (t-test, p=0.037) for the left temporal region in the gamma frequency band. (**B**) Preserved outward direction (i.e., directed towards other regions of the network) in the older group compared to the younger for the left temporal region, in the gamma frequency band. (**C**) Positive association between the increased of multiplex participation coefficient in inward direction in left temporal region in gamma1 frequency band and MMSE test (r= 0.327, p=0.017) in the older group. (**D**) Distribution of the young adults and older adults' subgroups for the multiplex participation coefficient in the left temporal region in gamma frequency band for the measure of directionality. (**E**) Positive association for the level of participation of left temporal region and VSTM score for the Low participation older subgroup (regression test, r= 0.485, p=0.035; no association with cognition for the High participation older subgroup). All results were adjusted for multiple comparisons using FDR corrections at $q < 0.05$
*p<0.05; **p<0.01; ***p<0.001

**Figure 3s: Balance of connectivity between the three layers in older group**

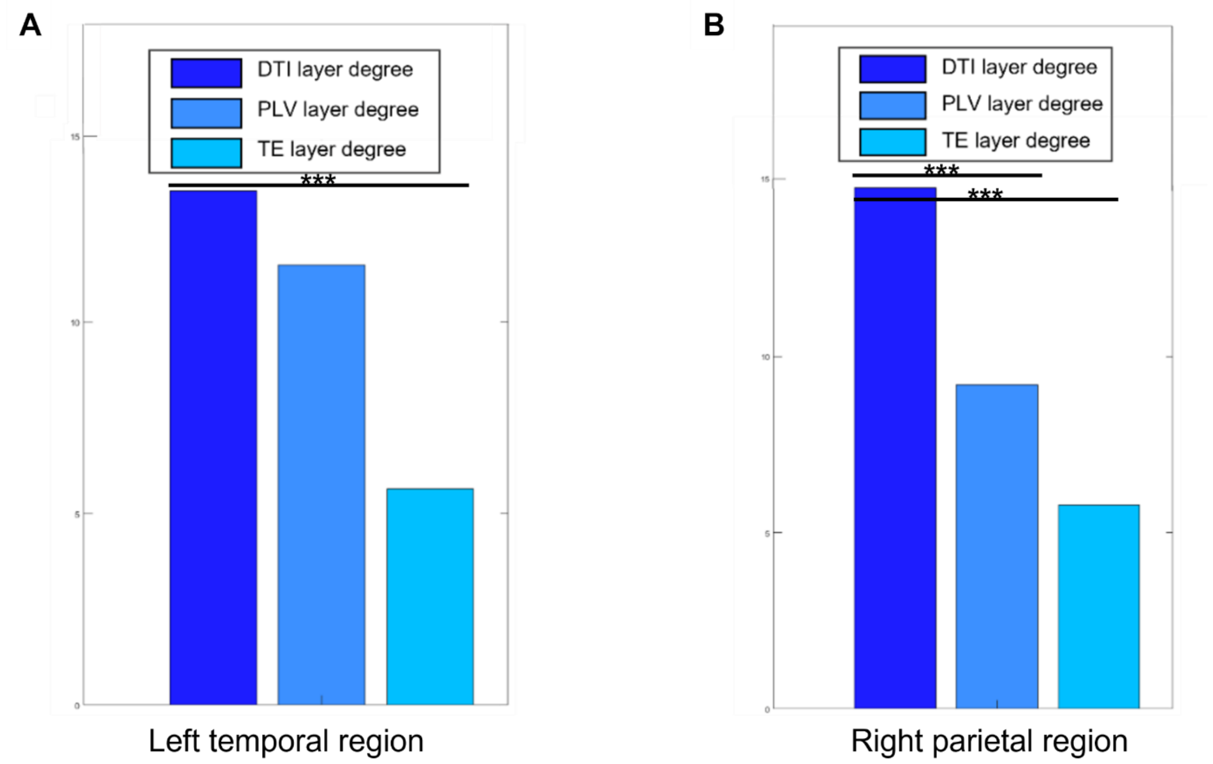


**Figure 3s**. (**A**) Balance of connectivity in the tree layers (DTI, PLV and TE) in the left temporal region in alpha frequency band in the older group. In dark blue, the degree of the DTI layer, in midlightblue, the degree of PLV, in light blue, the degree of the TE layer. Increased of DTI degree than TE degree (p<0.001). (**B**) Balance of connectivity in the tree layers (DTI, PLV and TE) in the right parietal region in alpha frequency band in the older group. In dark blue, the degree of the DTI layer, in midlightblue, the degree of PLV, in light blue, the degree of the TE layer. Increased of DTI degree than PLV and TE degree (p<0.001).
***p<0.001

**Figure 4s: Respective contribution of the three layers in younger group and both older subgroups**

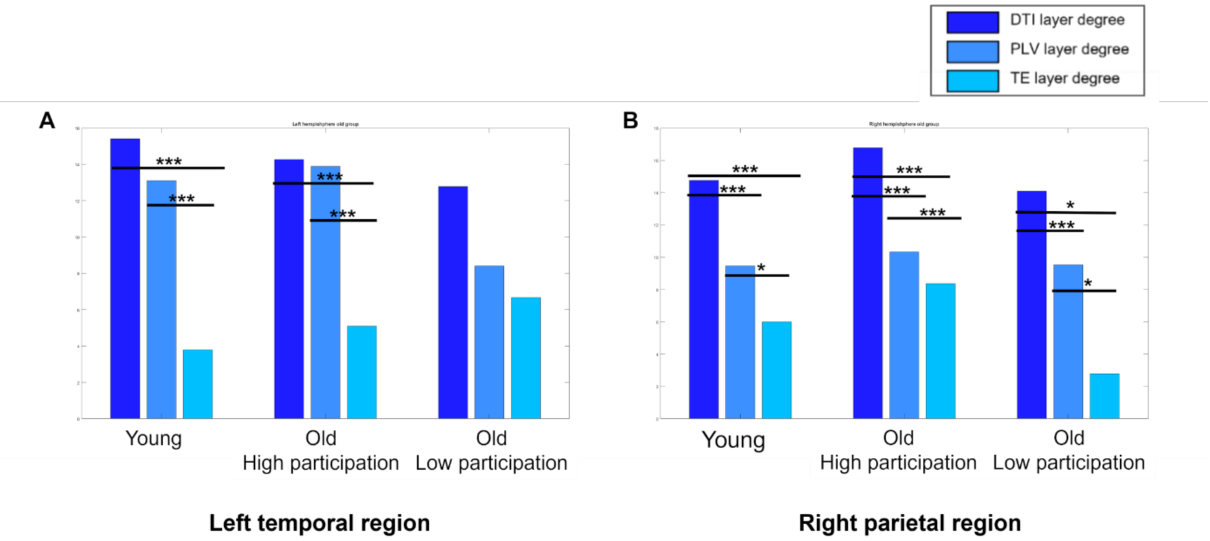


**Figure 4s.** (**A**) Balance of connectivity in the tree layers (DTI, PLV and TE) in the left temporal region in alpha frequency band in respectively in the younger group, High participation older group and Low participation older group. In dark blue, the degree of the DTI layer, in midlightblue, the degree of PLV, in light blue, the degree of the TE layer. (**B**) Balance of connectivity in the tree layers (DTI, PLV and TE) in the right parietal region in alpha frequency band in in respectively in the younger group, High participation older group and Low participation older group. In dark blue, the degree of the DTI layer, in midlightblue, the degree of PLV, in light blue, the degree of the TE layer.
